# vOMIX-MEGA: An ultra-fast end-to-end pipeline for terabyte-scale viral metagenomics analysis

**DOI:** 10.64898/2026.07.28.741255

**Authors:** Erfan Shekarriz, Elsa Vijendran, Joshua WK Ho

## Abstract

Viral identification for terabyte-scale metagenomic data is limited by scalability and computational resources. We present vOMIX-MEGA, an end-to-end viral metagenomic framework that overcomes performance bottlenecks by significantly improving parallelization and memory usage in critical steps. Benchmarked on empirical datasets, it completes processing in up to **50 minutes** with **24 GB of RAM**, bypassing four other state-of-the-art pipelines that require 7 hours (383 GB) to 14 days (32 GB). vOMIX-MEGA is on average 21% and 13% more accurate when benchmarked on mock and experimental data and is available via https://github.com/holab-hku/vOMIX-MEGA.

## 1. MAIN

Viruses are mobile genetic elements driving global biogeochemical cycles, ecological dynamics, and disease transmission across all domains of life [1], [2], [3]. Culture-independent viral metagenomics is used to explore a vast reservoir of viral dark matter [4]. Rapid advances in sequencing depth have made terabyte-scale datasets a routine standard in genomics research, generating a data surge that exposes severe bottlenecks in current computational workflows [5].

Established viral metagenomic software pipelines are fragmented and scale poorly, presenting an impractical trade-off: they either exhibit massive memory bloats requiring high-performance computing clusters or introduce extreme runtime latencies extending over weeks [6], [7]. Complicated deployment processes and software dependencies create further technical friction, restricting high-throughput analyses and undermining cross-study reproducibility [8]. This removes accessibility to large-scale viral metagenomic analysis for most researchers. To resolve these bottlenecks, we introduce vOMIX-MEGA, a containerized, end-to-end analytical pipeline engineered for high-performance, resource-accessible viral metagenomic profiling for terabyte-scale data.

vOMIX-MEGA’s efficiency is achieved by modifying core algorithms within standard workflows to resolve single-threaded dependencies and memory-intensive operations. We identified the evaluation of contig completeness and contamination via native CheckV [9], the only widely used tool for this purpose, as a primary performance bottleneck, as its RAM consumption scales exponentially with core provisioning (Supplementary Table 1). On terabyte-scale inputs, native CheckV requires up to 939 GB of RAM when utilizing 64 CPUs, an architecture inaccessible to most researchers (Figure 2a).

**Figure 1.**
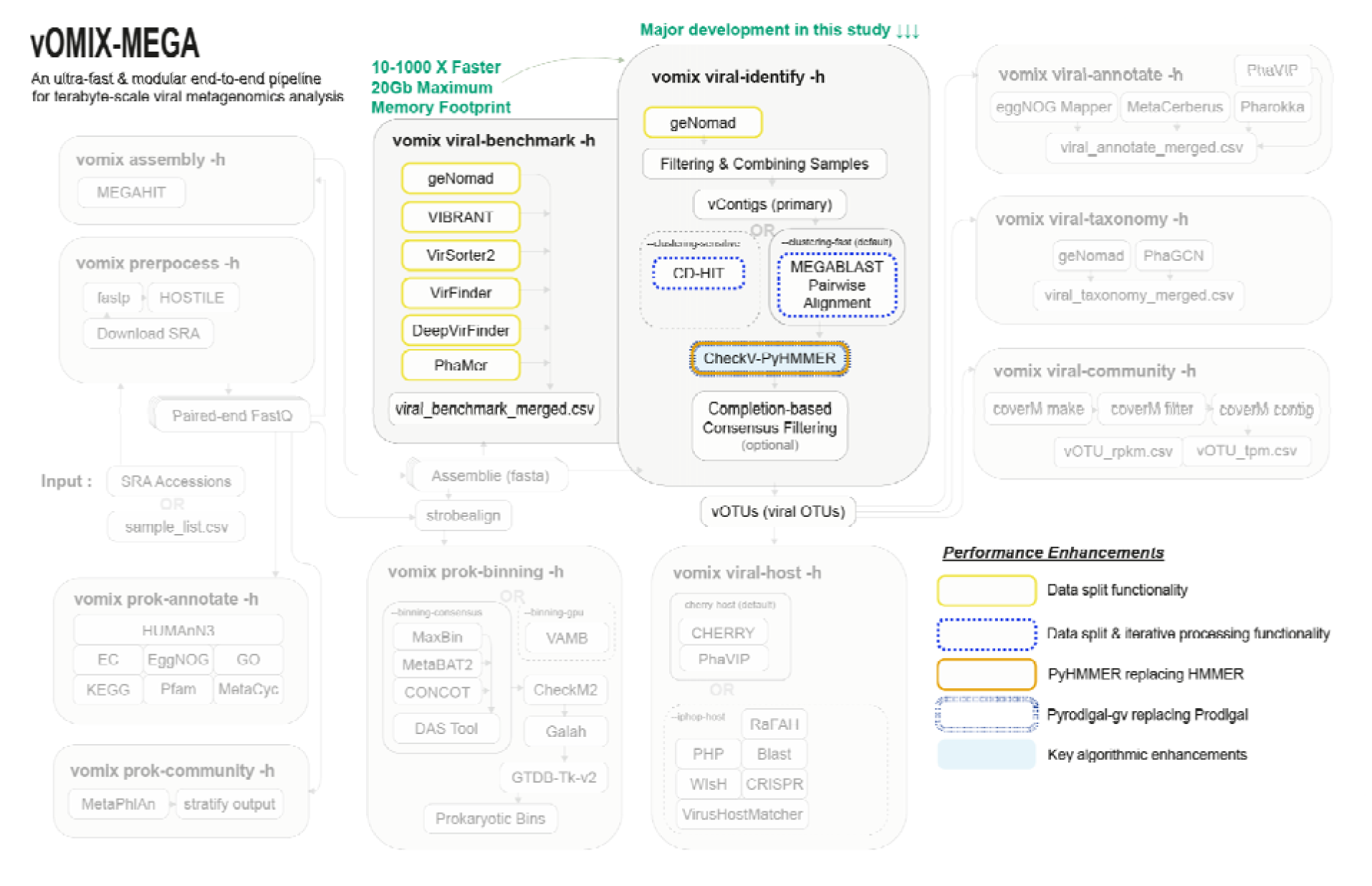
**Full pipeline overview of vOMIX-MEGA and underlying tools and modules**. Highlighted boxes include novel re-engineered algorithms that give vOMIX-MEGA its ability to process terabyte-scale data rapidly without memory bloats. Key performance and algorithm enhancements are marked accordingly.

**Figure 2.**
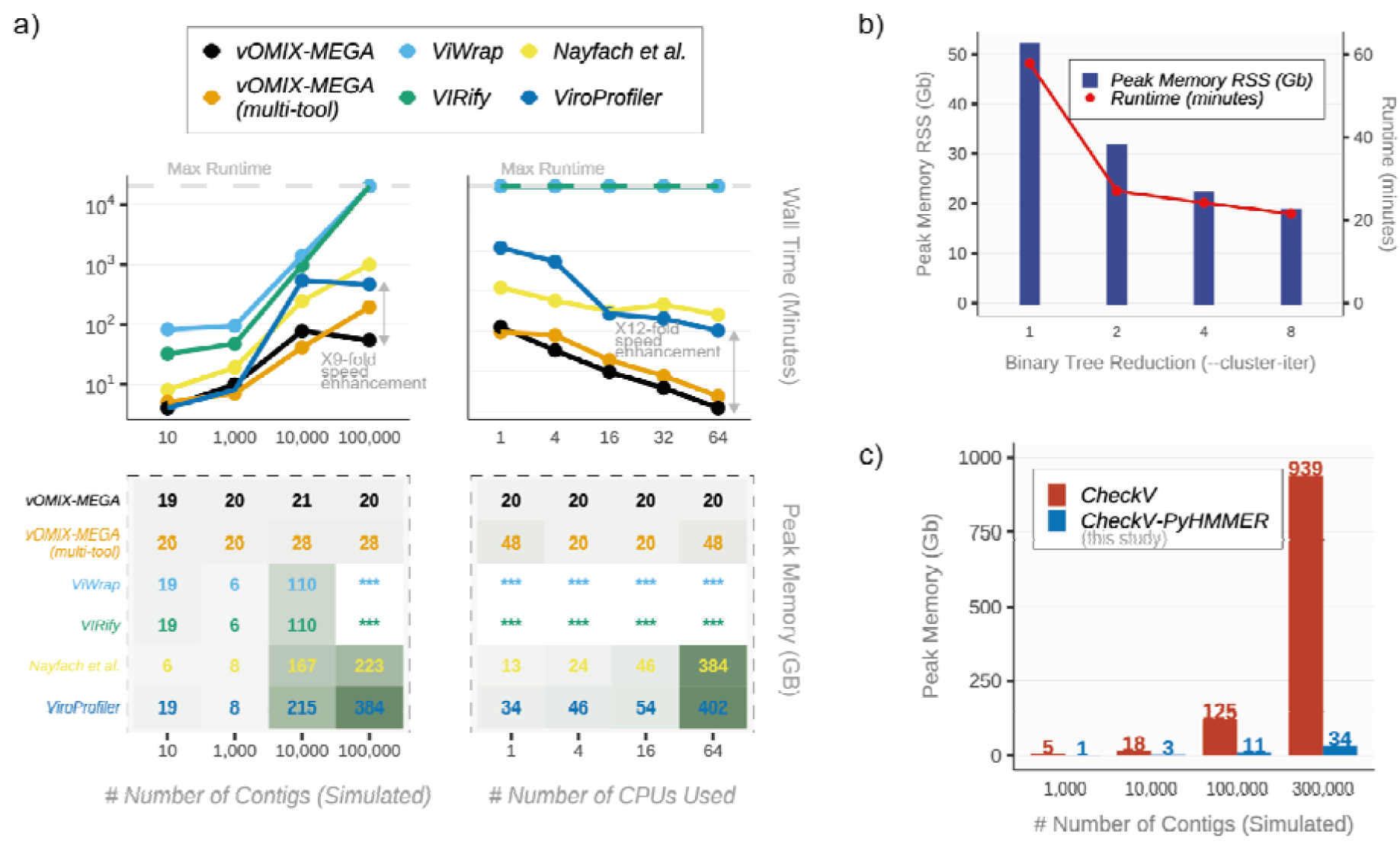
Scalability and runtime analysis of vOMIX-MEGA and other viral metagenomic pipelines. (a) Runtime and peak memory use analysis comparing vOMIX-MEGA, vOMIX-MEGA (multi-tool), ViWrap, VIRify, ViroProfiler, and the Nayfach et al. viral metagenomic pipelines. Analysis was carried out by using 64 CPUS across 10, 1000, 10000, and 100000 sequences, as well as using 1,4,16,32,64 CPUs with 50,000 sequences. Boxes marked with (***) mean that the run did not finish within the 14 days maximum runtime set for our analysis (b) Runtime and memory use of the different clustering methods including binary tree reduction (or tournament-style merger) reduction approach with --cluster-iter 1 (original), --cluster-iter 2, -- cluster-iter 4, and --cluster-iter 8 on 300,000 simulated contigs (c) Peak memory use of CheckV vs. CheckV-PyHMMER using 64 CPUs across an increasing number of contigs.

We bypassed this limitation by developing our own algorithmically revised version of CheckV called **CheckV-PyHMMER**, which accelerates quality assessment while maintaining annotation nearly identical to the original tool for all viruses (the differences being due to pyrodigal-gv’s higher sensitivity to giant viruses [10]) (Supplementary Table 2 and 3). CheckV-PyHMMER implements three structural optimizations: first, it decouples CPU-intensive contamination and completeness modules from the memory-heavy reference database module; second, it replaces native single-threaded Prodigal with Pyrodigal-gv [10], [11], which uses OpenMP parallelization optimized for alternative genetic codes and giant viruses; third, it substitutes standard HMMER with multi-threaded PyHMMER [12] Python bindings to achieve linear performance scaling (Supplementary Figure 1).

For memory-constrained local nodes and all viral identification tools, an optional data-splitting module (‘--contig-splits’, ‘--checkv-splits’) partitions large nucleotide fasta inputs into parallel fractions for isolated processing before performing a lossless concatenation, significantly lowering peak resident set size (RSS) (Supplementary Figure 2). Standard sequence-clustering algorithms (such as CD-HIT or CheckV’s native MEGABLAST-based pairwise alignment) likewise require constructing exhaustive sequence databases or performing all-versus-all alignments. The memory footprint (Resident Set Size; RSS) of these indexing and comparison steps scales quadratically with the number of input sequences. When scaling to terabyte-scale metagenomic datasets, this leads to massive memory inflation and execution hangs, frequently requiring computer systems with large memory that are inaccessible to most researchers. To bypass these hardware limitations, vOMIX-MEGA introduces an algorithmically re-engineered Divide and Conquer algorithm with a data-splitting and iterative clustering framework (‘--cluster-iter’). This module processes large nucleotide FASTA inputs in parallel, isolated fractions before performing hierarchical pooling and clustering, (Supplementary Figure 3), which we call a binary tree reduction (or tournament-style merger) reduction approach (See Methods for full introduction to our method). Depending on layers of clustering, our algorithm reduced the speed of clustering by CheckV’s ultrafast MEGABLAST method by up to 63%, and its peak memory usage (RSS) by up to 64% respectively (Figure 2b) (Supplementary Table 4). We benchmarked our approach using 1 (original algorithm), 2, 4, and 8 clustering iterations with both CD-HIT and MEGABLAST approach on the same set of contigs (Supplementary Table 12), and found a 93% identity in clustering sequencing representatives (final clusters) between all clustering iterations. (Supplementary Figure 6).

Finally and in general, we prevent memory-induced failures during large-scale operations by systematically prioritizing tools with stable memory footprints across vOMIX-MEGA. For example, ‘vomix viral-identify’ uses geNomad [10] as its core identification framework, omitting alternatives like DeepVirFinder [13] that exceed 300 GB RAM allocations at high thread counts (Supplementary Table 5).

We performed high-throughput scalability and runtime analysis under peak operational workloads, and benchmarked vOMIX-MEGA against four pipelines (ViroProfiler [14], Nayfach et al. [15], VIRify [16], and ViWrap [17]) alongside an internal vOMIX-MEGA (multi-tool) benchmark module using terabyte-scale datasets under a uniform allocation of 64 CPU cores (Fig. 2a). At n=100,000 contigs, default vOMIX-MEGA completed end-to-end profiling in **54 minutes** with a stable peak memory footprint of **20.1 GB**. Conversely, the non-optimized multi-tool module took 195 minutes and consumed 28.2 GB RAM due to DeepVirFinder’s thread overhead. Other standalone pipelines took substantially longer to analyze the same data and used significantly higher memory; ViroProfiler and Nayfach et al. resolved the cohort but required 464 minutes (384 GB RAM) and 998 minutes (223 GB RAM), respectively, bottlenecked by native CheckV execution. VIRify and ViWrap failed to complete processing within a strict 14-day tracking ceiling. This performance gap was preserved at lower sequence counts; at n=50,000 contigs, vOMIX-MEGA completed processing in 78 minutes (19.8 GB RAM), outperforming the mutl-tool (118 min, 29.1 GB RAM), Nayfach et al. (577 min, 189 GB RAM), and ViroProfiler (714 min, 354 GB RAM) (Supplementary Table 6).

Thread-scaling experiments on the n=50,000 contig tier (1,4,16,32, and 64 cores) demonstrated the parallel efficiency of vOMIX-MEGA (Figure 2a). Single-threaded execution required 357 minutes with a 19.7 GB RSS memory footprint; scaling to 64 cores monotonically reduced the runtime to 35 minutes while peak physical memory remained flat between 19.7 and 20.2 GB. The vOMIX-MEGA (multi-tool) identified all contigs in 308 to 49 minutes, but its memory expanded to 48.2 GB at 64 CPUs due to DeepVirFinder. Competing workflows displayed severe thread-dependent memory inflation: Nayfach et al. reached 384 GB RAM at 64 cores, and ViroProfiler spiked to 402 GB RAM. VIRify and ViWrap remained unresolved for 14 days, with core parallelization inflating peak physical memory to 136.2 GB and 148.8 GB, respectively (Supplementary Table 6).

Standalone stress tests supported the performance advantages of CheckV-PyHMMER over native CheckV at 64 CPUs (Figure 2c). At baseline volumes (n=1,000 and n=10,000), native CheckV displayed lower runtimes (58 s and 196 s) than CheckV-PyHMMER (192 s and 861 s) due to Python binding initialization overheads, but it demanded immense virtual memory (572.6–615.7 GB) compared to CheckV-PyHMMER (2.9–4.2 GB). At high-throughput limits (n≥100,000), CheckV-PyHMMER outpaced native execution. At n=100,000 sequences, CheckV-PyHMMER completed execution in 47.5 minutes using 11.0 GB RAM, while native CheckV required 53.5 minutes and 60.7 GB RAM. At n=300,000 contigs, native CheckV hit a severe processing wall, requiring 72.0 hours of continuous execution and an un-scalable 939.1 GB of physical RAM (1.31 TB virtual memory). Conversely, CheckV-PyHMMER completed the same analysis in 19.2 hours, marking a 3.7-fold reduction in runtime, while compressing peak physical RAM to 34.2 GB (a 27.4-fold memory reduction) and capping virtual memory at 36.1 GB. When benchmarked on a large dataset of 300,000 contigs derived from an expanded dataset of 41TB of data (See Methods), CheckV-PyHMMER was able to reduce runtime by 27% and 38%, and peak memory RSS by 21% and 46% when run with ‘--checkv-splits 3’ and ‘--checkv-splits 7’ respectively when compared to no splitting (‘--checkv-splits 0’) (Supplementary Table 7). With the --checkv-splits 7, CheckV-PyHMMER was able to maintain a maximum 18 GB memory footprint running this large dataset (Supplementary Table 7).

Our accuracy benchmarking on experimental and mock datasets was evaluated against a custom mock validation cohort and a multi-biome experimental dataset from Wu et al. [7] to isolate identification evaluation metrics across key tools (geNomad [10], DeepVirFinder [13], PhaMer [18], VirSorter2 [19], VirFinder [20], VIBRANT [21], PPR-Meta [22], Seeker [23]) (Supplementary Figure 4). The default engine of vOMIX-MEGA, geNomad, performed the best, reaching a Balanced Accuracy (BA) of 0.9785 on mock sequences and 0.8702 on experimental data. Alternative workflows followed: VIRify (0.8985 mock, 0.7845 experimental), PPR-Meta (0.8784 mock, 0.8259 experimental), ViWrap (0.8380 mock, 0.8296 experimental), and ViroProfiler (0.8380 mock, 0.8251 experimental). The Nayfach et al. pipeline demonstrated a low BA of 0.5041 (mock) and 0.5377 (experimental), nearing random classification (0.50). This suggests that the metagenomic gut virus compendium, compiled via this pipeline, may retain considerable host contamination despite structural size-filtering. Furthermore, consensus algorithms reduced taxonomic accuracy; the vOMIX-MEGA (multi-tool) voting scheme dropped BA to 0.7010 (mock) and 0.6189 (experimental), showing that a single optimized machine-learning classifier provides superior resolution while avoiding latency introduced by using multiple software (Supplementary Figure 5).

Our benchmarking revealed a specificity-sensitivity imbalance across alternative tools (Supplementary Figure 5). While most platforms exhibited high sensitivity for authentic viral contigs, exemplified by the vOMIX-MEGA (multi-tool) approach reaching 0.9918 (mock) and 0.9950 (experimental), they accumulated higher false-positive rates due to poor specificity. On the mock cohort, specificity dropped drastically for VirSorter2 (0.5728), the multi-tool approach (0.4101), VirFinder (0.2631), and VIBRANT (0.0843). This pattern was also observed with experimental data benchmarking, where VirSorter2 and VIBRANT logged specificities of 0.3805 and 0.3630. The Nayfach et al. workflow coupled low sensitivity (0.6270 mock, 0.7102 experimental) with low specificity (0.3811 mock, 0.3653 experimental), introducing substantial background noise that could mislead downstream metabolic and ecological annotations. Conversely, default vOMIX-MEGA maintained precise, balanced discrimination, delivering high specificities (0.9826 mock, 0.8104 experimental) alongside robust sensitivity thresholds (0.9743 and 0.9300), offering a reliable computational end-to-end pipeline for standardized viral metagenomic discovery (Supplementary Table 8 and 9).

These results demonstrate that vOMIX-MEGA offers a standardized, resource-efficient solution for terabyte-scale viral metagenomic data analysis.

## 2. ONLINE METHODS

### 2.1 Pipeline Architecture and Implementation

The vOMIX-MEGA framework is implemented as an automated, reproducible workflow built on a Snakemake [24] back-end and is fully containerized using Docker and Singularity/Apptainer to ensure cross-platform environment stability. Detailed methods regarding software compilation, algorithmic re-engineering of core back-end components (such as the integration of CheckV-PyHMMER and Pyrodigal-gv), and optimization protocols are described extensively in the Supplementary Methods section of this study.

### 2.2. Runtime Analysis and Scalability Benchmarking

The development of vOMIX-MEGA was prompted by critical computational bottlenecks and memory limitations observed during the baseline evaluation of a real-world hypertension patient gut metagenomic contig collection comprising 41,853 contigs across 420 samples. To systematically assess pipeline scalability under high-throughput processing loads, this baseline repository was integrated into a larger evaluation framework alongside three independent, terabyte-scale empirical datasets together making the following:

- Cow gut microbiome dataset [25]: 0.8 TB, 9,843 initial contigs.
- Mouse gut metagenome dataset [26]: 8.2 TB, 48,674 initial contigs.
- Hypertension human gut microbiome dataset [27]: 41 TB, 41,853 initial contigs.
- Global gut microbiome dataset: 60 TB, 126,789 initial contigs (Internal data gathered for this study)

To eliminate database-size confounding variables and enable clear cross-pipeline visualization, these empirical sequences were converted into six standardized benchmarking cohorts. Target sequence thresholds were achieved by applying random subsampling (without replacement) to reduce sequence volume, or random duplicate sequence injection to expand volume where appropriate. This standardization protocol generated six final validation datasets scaling precisely to n = 10, 1,000, 10,000, 50,000, 100,000, and 300,000 contigs.

Comparative performance profiles were executed across four alternative viral metagenomic workflows, ViroProfiler, VIRify, ViWrap, and the Nayfach et al. pipeline, as well as the default vOMIX-MEGA pipeline and its standalone vOMIX (multi-tool) module. While the default vOMIX-MEGA framework relies on optimized sequence processing dependencies, the multi-tool benchmark module sequentially performs non-optimized tools (geNomad, DeepVirFinder, PhaMer, VirSorter2, VirFinder, and VIBRANT) within a single environment to provide users with direct comparative controls. For workflows where automated viral quality assessment was not natively supported (the Nayfach et al. pipeline and ViroProfiler), a standalone implementation of native CheckV was manually integrated into the execution stream to ensure uniform downstream evaluation of contig contamination and completeness metrics.

All benchmarking instances were deployed on identical, isolated high-performance computing nodes across increasing thread allocation (1, 4, 16, 32, and 64 CPU cores). Hardware resource allocation and execution efficiency were systematically noted using Snakemake’s default benchmarking capability. The framework continuously monitored and documented five primary computational dimensions:

- **Walltime**: Total elapsed execution duration from data ingestion to output delivery, quantified in minutes.
- **Memory Utilization**: Multidimensional RAM consumption profiles tracking Virtual Memory Size (VMS), Resident Set Size (RSS), Unique Set Size (USS), and Proportional Set Size (PSS).
- **Data Throughput**: Exact input and output file storage sizes.
- **Mean Load**: The average computational core utilization throughout active execution windows.
- **CPU Time**: Cumulative processing time utilized by allocated hardware threads.

A maximum execution ceiling was strictly enforced at 720 hours per individual pipeline run. Any analysis that failed to complete its assigned sequence tier within this allocated window was terminated and logged as unresolved.

### 2.3 Data splitting and iterative clustering algorithm

The iterative clustering algorithm is structured as a binary tree reduction (or tournament-style merger) (Supplementary Figure 3). The depth and branching factor of the tree are dynamically controlled by a user-defined parameter ***L***, representing the total clustering iterations (‘--clustering-iter’).

The system operates according to the following mathematical rules:

- **Initial Partitioning**: The raw input FASTA is divided into ***N*_start_** starting chunks: ***N*_start_ = 2^L−1^**
- **Layer and Chunk Indexes**: Let ***l*** denote the current layer of the reduction tree, where ***l* ∊ {1, 2,…, *L*}**. Let ***c*** denote the chunk index within layer ***l***, where ***c* ∊ {0, 1,…, 2^L−1^ − 1}**.
- **Base split inputs**. Let ***S_c_*** represent the ***c-***th partitioned raw FASTA file.
- **Clustering Operator**: Let **Cluster(*X*)** denote the sequence clustering and dereplication function, executed using either CD-HIT or CheckV’s MEGABLAST-based pairwise alignment.

The recursive generation of clustered outputs ***C_i,c_*** at any given node of the tree is defined by the recurrence relation.

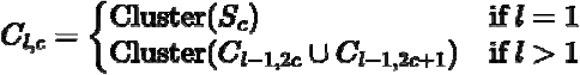

Using this binary tree architecture, the total number of individual clustering executions (***N*_total_**) required to resolve the complete dataset down to a single, fully non-redundant output chunk (***C_L_*_,0_**) is given by:

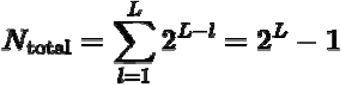

The formal mathematical proof of why this architecture is guaranteed to reduce memory reduction can be found in the **Supplementary Methods** of this paper.

○ Viral Identification Accuracy Benchmarking

All viral metagenomic pipelines and their underlying tools were benchmarked using a custom mock contig dataset alongside a recently published experimental viral benchmarking dataset developed by Wu et al. All validation sequences were analyzed using the complete alternative workflows described above, as well as the vOMIX-MEGA viral-benchmark module to isolate the standalone classification metrics of geNomad, DeepVirFinder, PhaMer, VirSorter2, VirFinder, VIBRANT, PPR-Meta, and Seeker.

Diagnostic criteria, including Sensitivity, Specificity, Balanced Accuracy, and Receiver Operating Characteristic (ROC) curves, were calculated using the R package caret [28]. The underlying performance parameters were defined mathematically using the following expressions:

- **Sensitivity** = TP / (TP + FN)
- **Specificity** = TN / (TN + FP)
- **Balanced Accuracy** = (Sensitivity + Specificity) / 2
- **F1-score** = (2 × TP) / ((2 × TP) + FP + FN)

where TP represents True Positives, TN represents True Negatives, FP represents False Positives, and FN represents False Negatives.

#### 2.3.1 Mock Benchmarking Cohort

To assemble the mock dataset, we downloaded all complete viral genomes (n = 4,143), complete bacterial genomes (n = 3,020), chromosomal/complete eukaryotic genomes (n = 5,600), and all plasmid sequences (n = 58,289) deposited to NCBI’s RefSeq database after July 21st, 2023. This cutoff date matches the public training freeze of geNomad to ensure an unbiased evaluation of the newest model architecture. Eukaryotic and plasmid sequences were included specifically to evaluate false-positive misclassification rates common among k-mer based approaches.

Following the approach outlined by Wu et al., we quantified the similarity of post-July 21st, 2023 viral genomes against older entries using MMseqs2 [29] (easy-search with translated type --search-type 2). Viruses were partitioned into three categories based on evolutionary divergence:

- Low identity: 20% similarity to older RefSeq entries.
- Medium identity: >20% and 40% similarity to older RefSeq entries.
- High identity: > 40% and 100% similarity to older RefSeq entries.

All genomes were fragmented into non-overlapping adjacent fragments of lengths 500, 1,000, 3,000, 5,000, and 10,000 nucleotides. From these pools, 10,000 sequences were selected at random for each primary taxonomic division (viruses, bacteria, eukaryotes, and plasmids). For prokaryotic and eukaryotic fragments, CheckV was applied to remove potential proviruses before combining all segments into a unified test dataset.

#### 2.3.2 Experimental Benchmarking Cohort

For empirical validation, fully annotated contigs were obtained directly from the multi-biome environmental database compiled by Wu et al. [30]

## 3. CODE AVAILABILITY

The **vOMIX-MEGA** package, source code, and installation instructions are publicly available on GitHub at https://github.com/holab-hku/vOMIX-MEGA, with comprehensive documentation hosted at https://vomix-mega.readthedocs.io/en/latest/. The underlying workflow engine, **vomix-snakemake**, is available at https://github.com/holab-hku/vomix-snakemake/wiki. All custom scripts used to generate the figures and analyses presented in this study are archived at https://github.com/erfanshekarriz/vomix-manuscript.

## 4. ACKNOWLEDGEMENTS

The authors gratefully acknowledge financial support from the Hong Kong PhD Fellowship Scheme (HKPFS) awarded by the Research Grants Council (RGC) of Hong Kong. We also extend our gratitude to the Li Ka Shing Faculty of Medicine and the School of Biomedical Sciences at The University of Hong Kong for providing institutional support and infrastructure.

## 5. AUTHOR CONTRIBUTIONS

**E.S.** conceptualized the project, developed the algorithms, performed data analysis and generation, and wrote the manuscript. **E.V.** contributed software code and designed the architecture of the command-line interface wrapper script. **J.W.K.H.** conceptualized the project, supervised the research and provided critical intellectual input and manuscript edits.

## 7. DATA AVAILABILITY

All data used to generate the figures and findings in this study are available at https://github.com/erfanshekarriz/vomix-manuscript.

## 9. SUPPLEMENTARY INFORMATION

Supplementary Information is available for this paper (comprising Supplementary Methods, Supplementary Figures, and Supplementary Tables).

