## Supplementary Figures for "vOMIX-MEGA: An ultra-fast end-to-end pipeline for terabyte-scale viral metagenomics analysis"

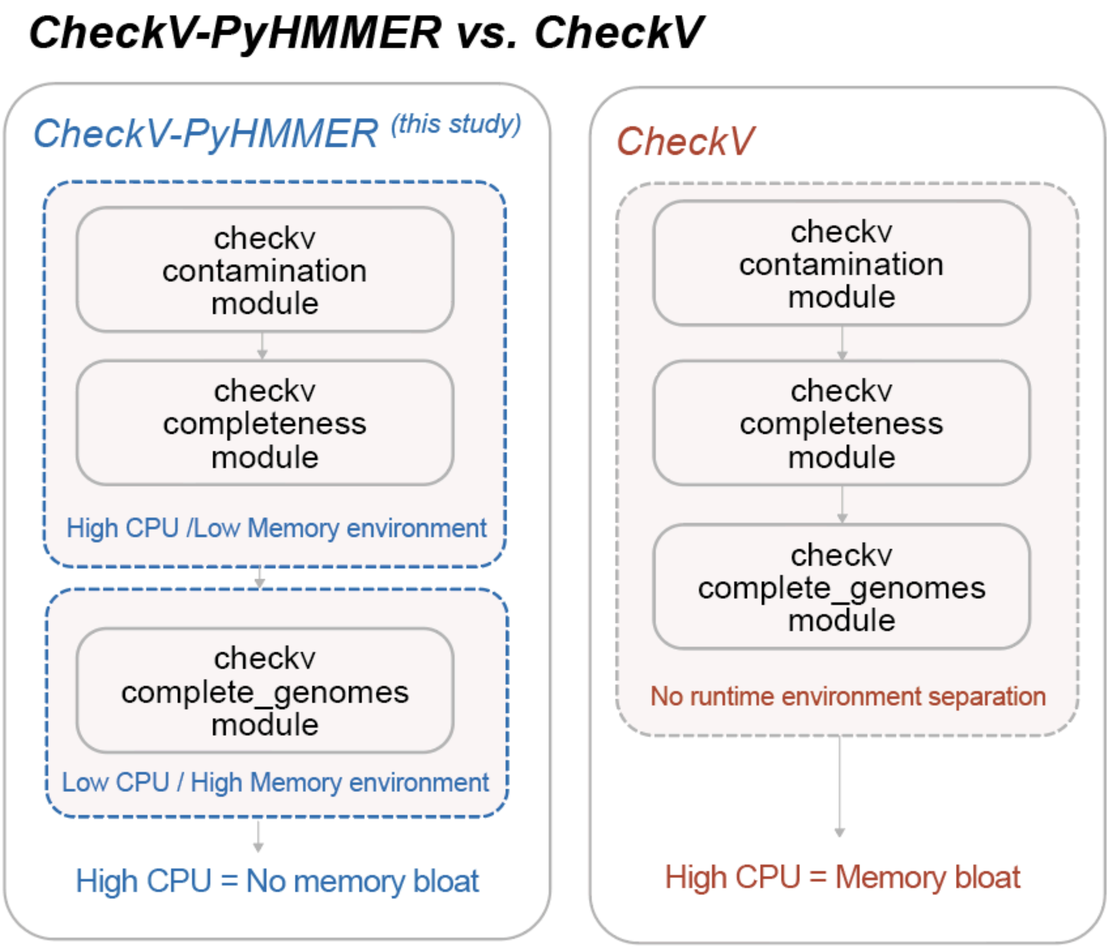


**Supplementary Figure 1** | Algorithm overview differences between CheckV vs. CheckV-PyHMMER (this study)

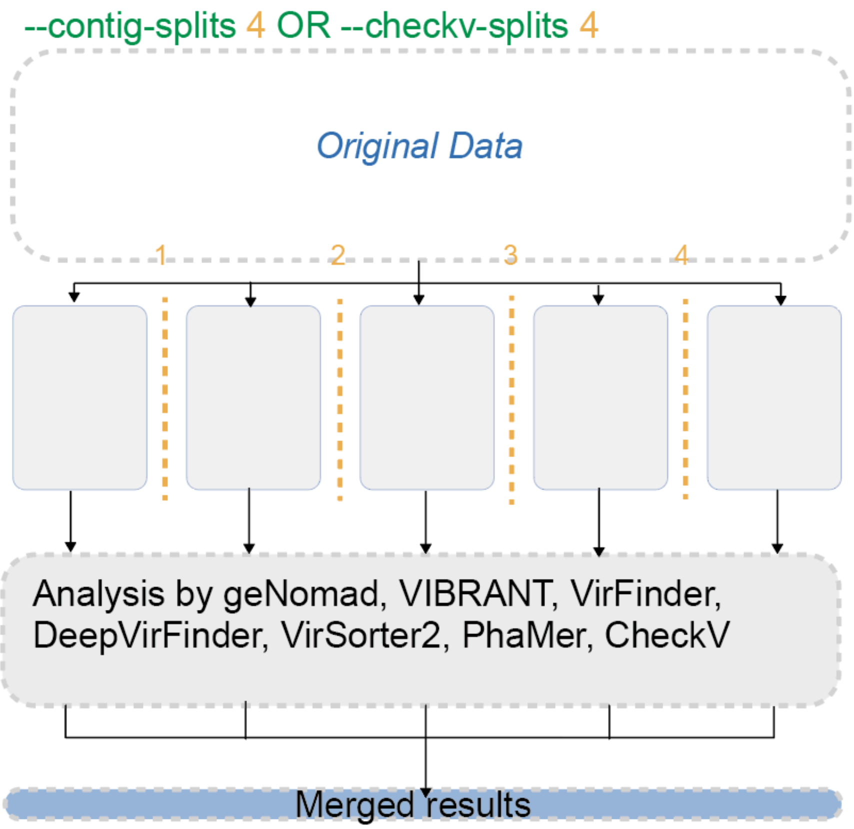


**Supplementary Figure 2 |** Figure showing data splitting functionality used in “checkv-split” (for checkv-pymer module) or “contig-split” (used in any module where applicable) which reduces RSS memory by parallelising processing where the functionality is not yet implemented.

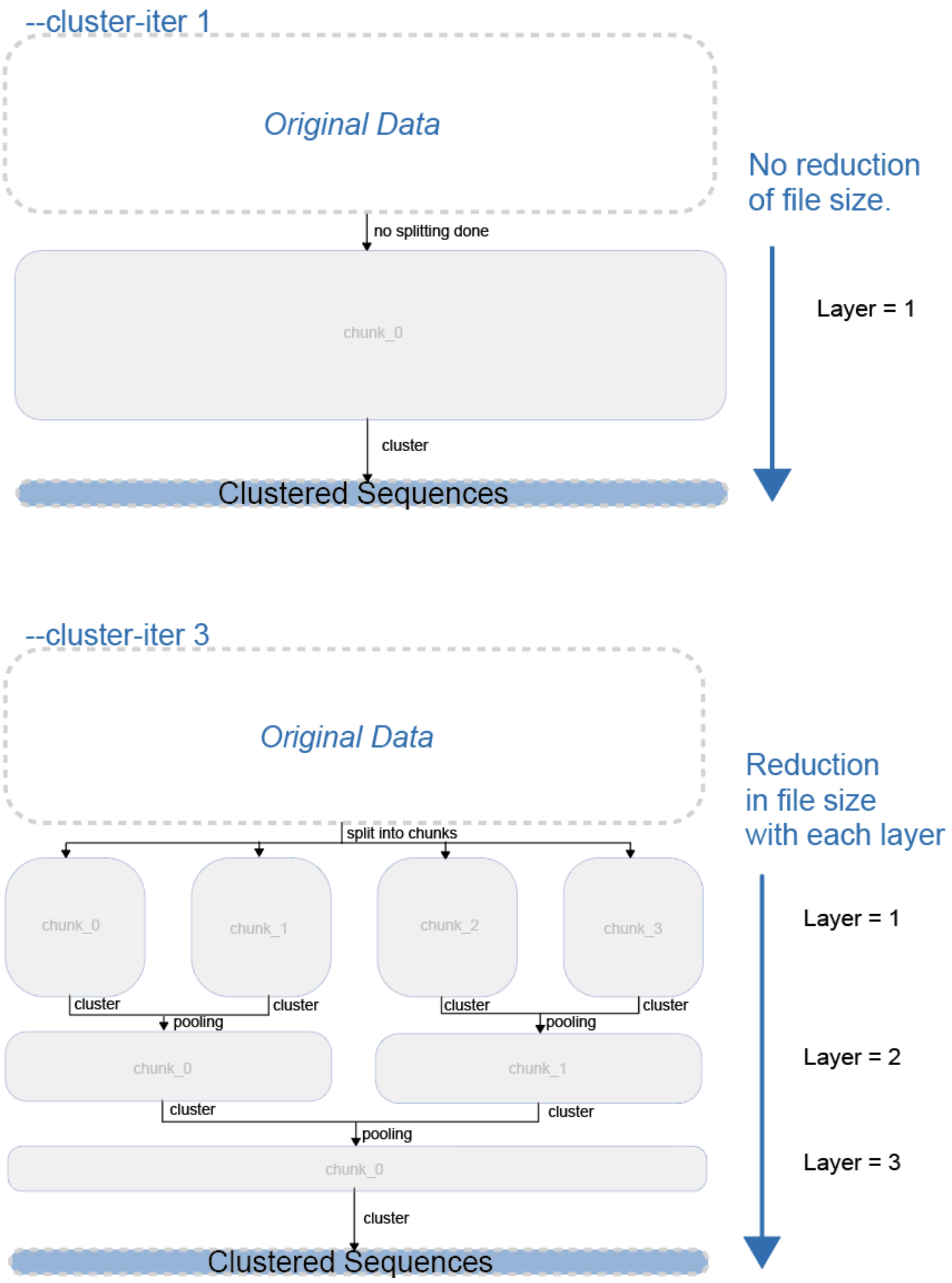
--cluster-iter 1
Original Data
no splitting done
chunk_0
cluster
Clustered Sequences
No reduction
of file size.
Layer = 1
--cluster-iter 3
Original Data
split into chunks
chunk_0
chunk_1
chunk_2
chunk_3
cluster
pooling
chunk_0
chunk_1
cluster
pooling
chunk_0
cluster
Clustered Sequences
Reduction
in file size
with each layer
Layer = 1
Layer = 2
Layer = 3

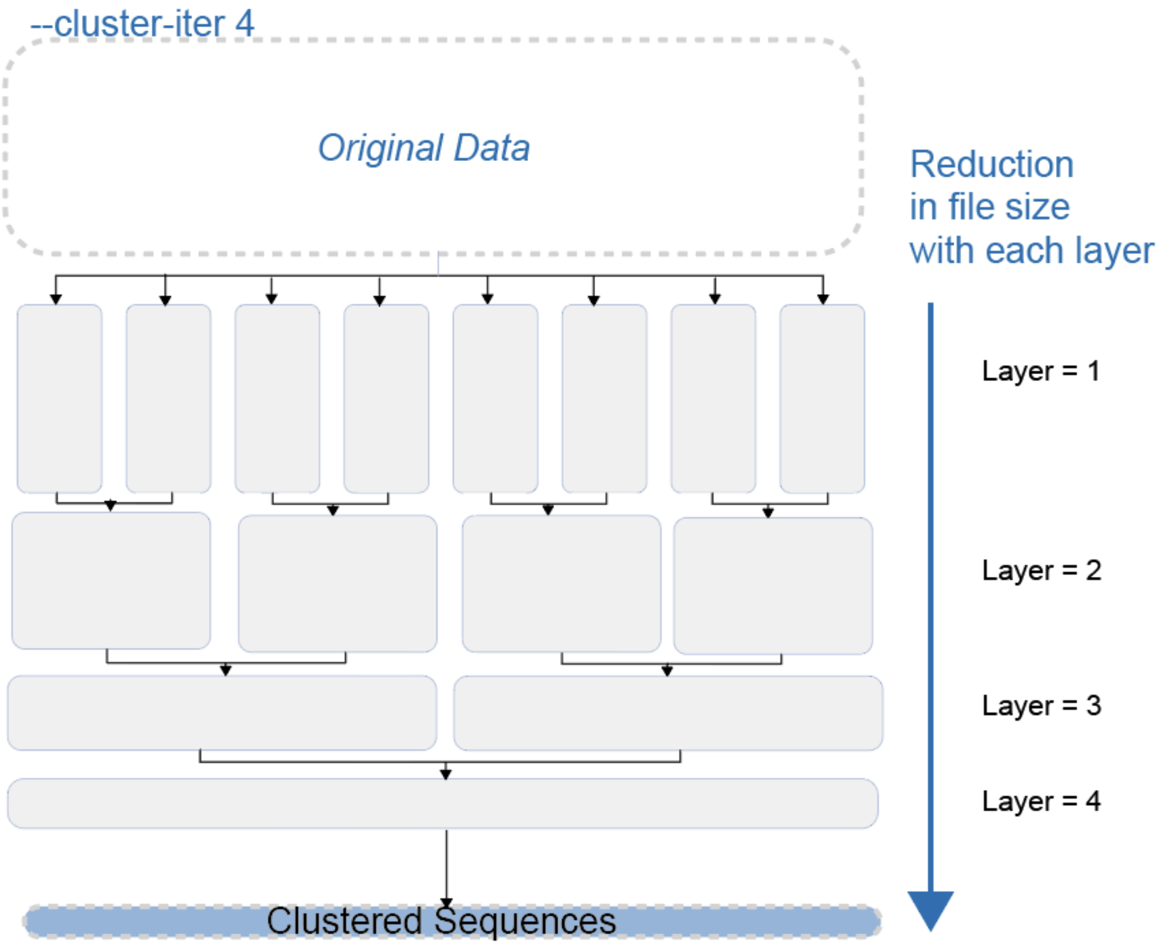


**Supplementary Figure 3** | Schematic diagram for the binary tree reduction (or tournament-style merger) reduction approach that splits the data into N layers (N=1 being no splitting of data equivalent to bruteforce algorithm) showing N=1, N=3, and N=4 splits (--cluster-iter 1, --cluster-iter 3, --cluster-iter 4).

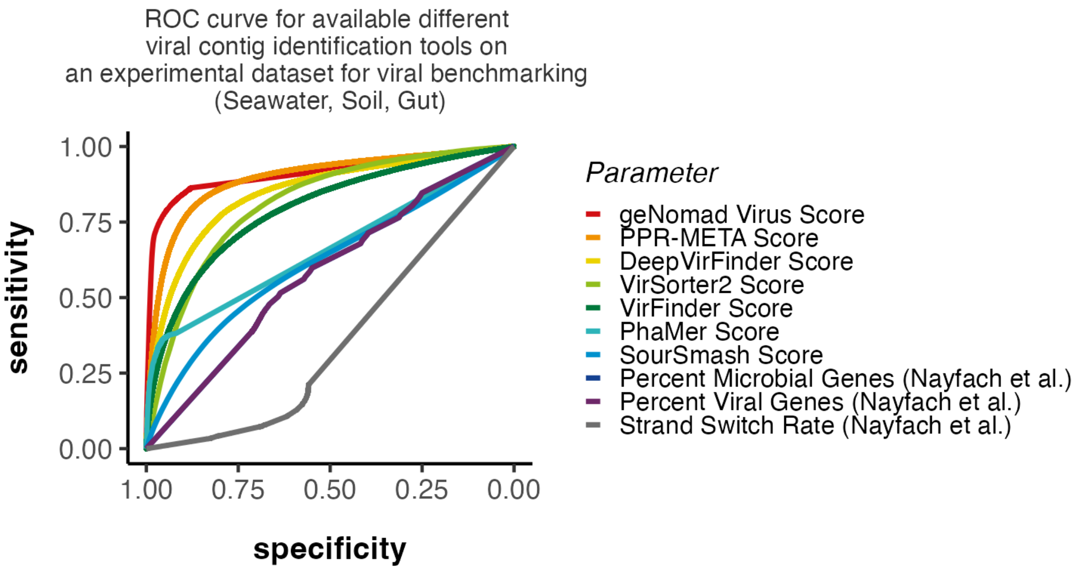


**Supplementary Figure 4** | Tool-based sensitivity vs. specificity analysis on experimental and mock viral contig datasets using the vomix viral-benchmark module.

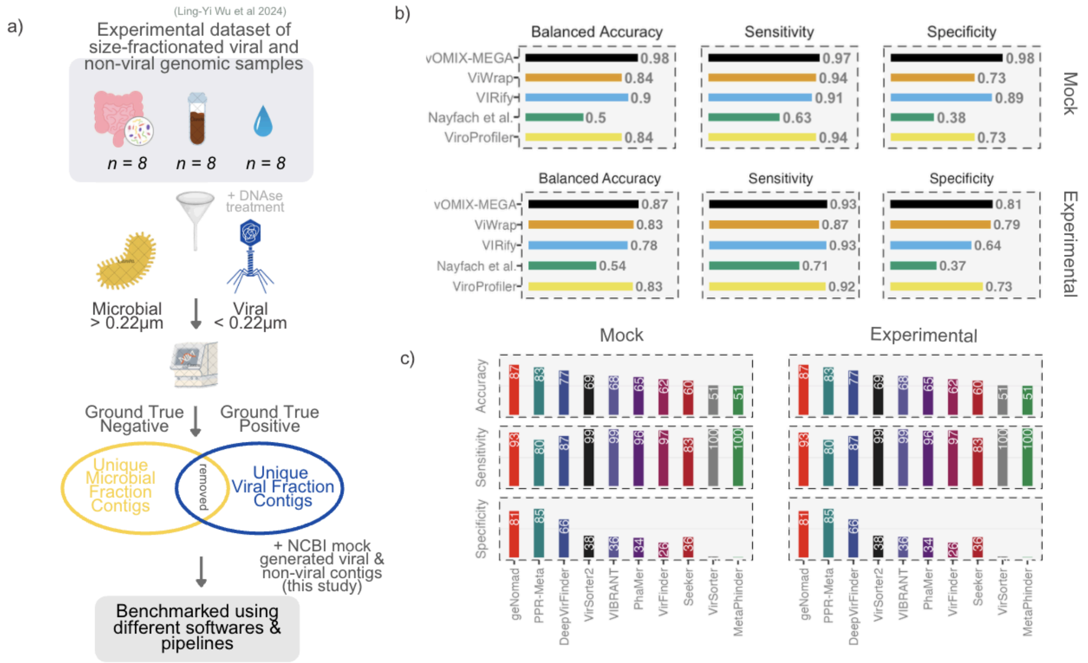


**Supplementary Figure 5** | Tool-based sensitivity vs. specificity analysis on experimental and mock viral contig datasets. (a) Shows the Wu Et Al [1] experimental benchmarking dataset schematic diagram. (b) Shows the pipe-line level and (c) shows the software-level benchmarking of experimental and mock datasets.

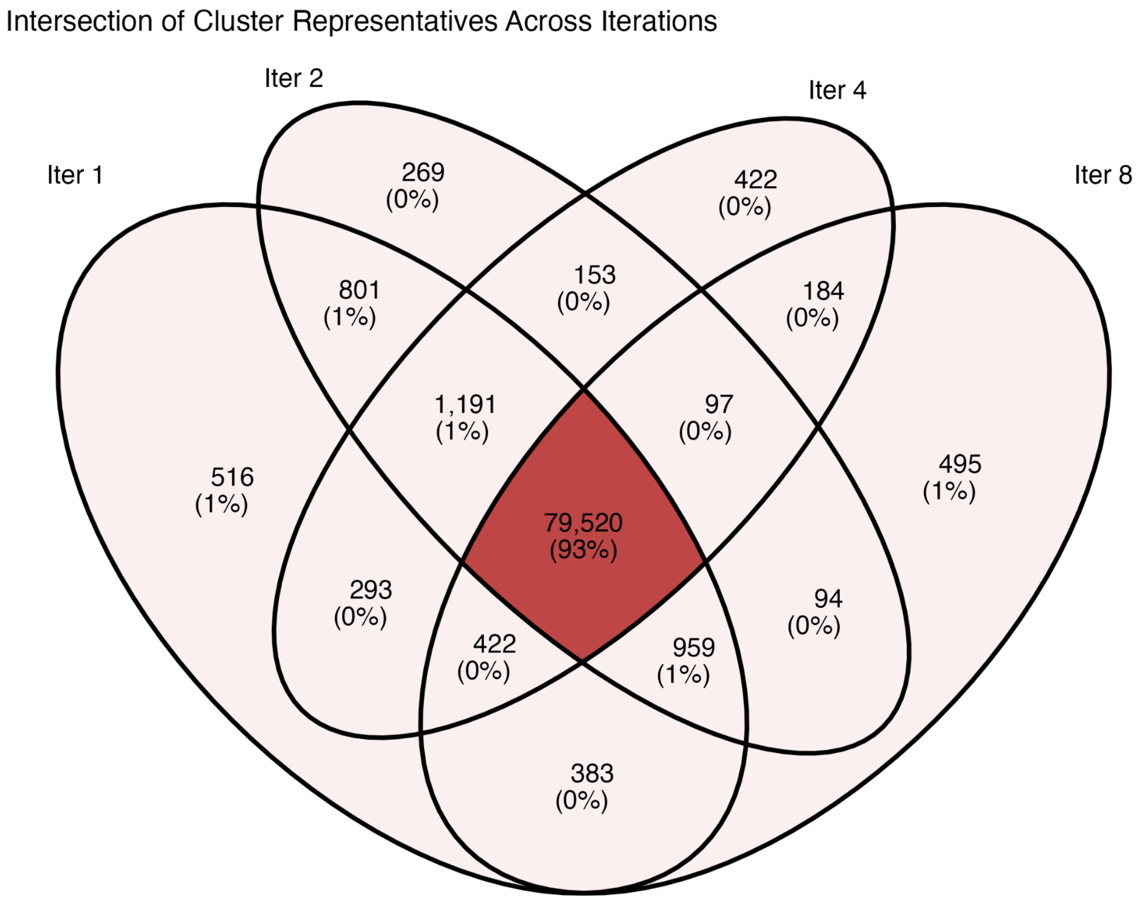


**Supplementary Figure 7 |** Clustering iteration identity test using 300,000 contigs and MEGABLAST approach. Venn diagram representing the number of unique sequence representatives (final clusters).
