## Supplementary Methods for "vOMIX-MEGA: An ultra-fast end-to-end pipeline for terabyte-scale viral metagenomics analysis"

### 1. SUPPLEMENTORY METHODS

#### 1.1. vOMIX-MEGA: A Comprehensive Modular Overview

vOMIX-MEGA is built on a Snakamake backbone and we use its schemas to structure modules and sort out jobs. Each module is in a Snakemake file (.smk) which will run via a wrapper script as input by the user via the command line (*vomix -h*). Having this backbone, each job can be individually modified, configured, and fine-tuned according to the needs of the user, allowing flexibility for different applications for the user. Each analytical task is logged, benchmarked, assigned to cluster if applicable, and sorted in a structured manner. Here, we will describe the different modules and softwares behind vOMIX-MEGA. Full documentation of modules and their outputs can be found at <https://github.com/holab-hku/vOMIX-MEGA/wiki> .

##### 1.1.1. vomix setup-database -h

vOMIX-MEGA handles all ad-hoc steps of viral metagenomics, including cumbersome database setups for each software. The `*vomix setup-database`* module includes the installation of all databases needed for full vOMIX-MEGA usage. However, when running any of the modules below, vOMIX-MEGA pre-determines each database needed for the module and automatically downloads it using the `--setup-database` flag. The size of each database and available free disk space is disclosed when running the dry run (`--dry-run` or `-n-`).

##### 1.1.2. vomix preprocess -h

The preprocess module requires a *sample_list.csv* input file (See Section 7.2) or alternatively a list of SRA accessions as input. If SRA accessions are given, each is automatically validated and downloaded using NCBI Datasets command-line tools [[1]](https://www.zotero.org/google-docs/?hoohl7). Locally stored paired-end gunzipped fastq files can also be provided as input. After downloading and validation, the fastq files are quality controlled using fastp [[2]](https://www.zotero.org/google-docs/?V9yaTV), and optionally host-decontaminated using Hostile [[3]](https://www.zotero.org/google-docs/?ycM5zO) (*--decontam-host*). By default, Hostile has multiple host databases that can be automatically downloaded, but users can make their own custom database of their unique host depending on experimental setup (Read more <https://github.com/bede/hostile>). MultiQC [[4]](https://www.zotero.org/google-docs/?KTf69e) then summarises the preprocessing analyses and generates an *html report*, which can be used to assess sample quality. The output of the preprocess module is a pair of cleaned, optionally decontaminated, fastq files used for downstream analysis.

##### 1.1.3. vomix assemble -h

The assembly module likewise requires a *sample_list.csv* input file with associated cleaned, decontaminated paired-end fastq files generated from *vomix preprocess* and assembles them using MEGAHIT [[5]](https://www.zotero.org/google-docs/?QTLuYi). Co-assembly is possible through modifying the *sample_list.csv* by setting the “assembly” column as the same for multiple Sample IDs. An assembler can be chosen using the --assembler MEGAHIT flag, as we are developing a MetaviralSPAdes [[6]](https://www.zotero.org/google-docs/?Pv3vOs) option, although co-assembly is currently not available for SPAdes. The output of this module is a .fasta contig file per sample, as well as a contig summary report showing the size and distribution of contigs per sample.

##### 1.1.4. vomix viral-identify -h

The *viral-identify* module is the core of vOMIX-MEGA’s developments on speed enhancement and memory efficiency. We have in parallel implemented a viral contig identification benchmarking module (*vomix viral benchmark*) that analyses contig files using 8 different viral identification tools (geNomad [[7]](https://www.zotero.org/google-docs/?mrEmzS), DeepVirFinder [[8]](https://www.zotero.org/google-docs/?MPYkIS), Phamer2 [[9]](https://www.zotero.org/google-docs/?lQvfaf), VirSorter2 [[10]](https://www.zotero.org/google-docs/?kOAnQB), VirFinder [[11]](https://www.zotero.org/google-docs/?V8gBhP), VIBRANT [[12]](https://www.zotero.org/google-docs/?y5WFXm), PPR-Meta [[13]](https://www.zotero.org/google-docs/?8AbqB2), and Seeker [[14]](https://www.zotero.org/google-docs/?JvRRCf)) and summarises their outputs in one file for the user to allow comparison.

The *viral-identify* module is built on the principle that more tools do not necessitate better results, as a principle of machine learning that there exists a trade-off between sensitivity and specificity [[15]](https://www.zotero.org/google-docs/?KcSAU3). Multi-tool approaches that fail to complement each other with new information gain, often lead to lowered sensitivity and hence cannot correctly identify most viral contigs ( Supplementary Figure 5). While consensus multi-tool approaches to identifying viral sequences have been widely adopted in the field of metagenomics [[16], [17], [18], [19]](https://www.zotero.org/google-docs/?41cKGL), we demonstrate using benchmarking that one optimized tool is sufficient for achieving the most accurate balance between sensitivity and specificity as seen when multi-tool pipelines have lower accuracies than individual tools on benchmarked data (Supplementary Figure 5)

The identification module first length-filters all sequences according to user input (--contig-minlen 0) and determines whether they are viral or non-viral using *geNomad*. According to our benchmarking, geNomad is the most accurate tool when benchmarked on both experimental and mock data (See Fig 3). geNomad was likewise selected as it has a stable memory footprint of ~ 20 ± 5 Gb across different CPUs used, and is scalable with increasing number of CPUs. The outputs of geNomad across different samples are then filtered according to user-specified thresholds (default 0.7), and combined into one file, generating pre-clustered viral contigs (vContigs). The vContigs are then clustered using CD-HIT (--clustering-sensitive) or CheckV’s fast MEGABLAST-based clustering algorithm (--clustering-fast) [[20]](https://www.zotero.org/google-docs/?Vd7fAT) with 95% average nucleotide identity and 85% coverage according to the Minimum Information about an Uncultivated Virus Genome (MIUViG) guidelines [[21]](https://www.zotero.org/google-docs/?5xFuzF). The quality of the clustered vContigs is then assessed in terms of contamination and completeness using CheckV-PyHMMER.

In this study we have developed CheckV-PyHMMER as a much faster and memory-efficient version of CheckV. Specifically, we have i) restructured CheckV’s processing algorithm to decouple CPU intensive `checkv *contamination`* and *`*checkv *completeness`* module from the memory intensive `checkv *complete_genomes`* module ii) replaced Prodigal [[22]](https://www.zotero.org/google-docs/?SbAy4G) with Pyrodigal-gv (Pyrodigal is parallelised version of Prodigal and Pyrodigal-gv <https://github.com/althonos/pyrodigal-gv> has a special database for The giant virus and alternative genetic code virus parameters), and iii) replaced non-scalable HMMER [[23]](https://www.zotero.org/google-docs/?JjMLZO) with a multi-thread PyHMMER [[24]](https://www.zotero.org/google-docs/?2nylYP) to allow scalability with increasing number of CPUs ( Supplementary Table 1). Overall CheckV-PyHMMER runs a faster, more scalable, and more memory-efficient analysis (See Fig 2c), while maintaining 99.9%> identical output as the original CheckV software (The reason for the negligible <0.1% difference in output is in that CheckV-PyHMMER uses prodigal-gv for protein annotation, while CheckV uses prodigal. The former is sensitive to giant viruses and hence can identify a small number of distinct proteins that prodigal cannot, resulting in different outputs. This does not change the underlying HMMER search algorithm which would otherwise produce identical outputs to the original CheckV) (Supplementary Figure 2 and 3) . If users wish, they may use the original CheckV, the may do so with --checkv-original flag. The CheckV-PyHMMER is a main reason for why vOMIX-MEGA’s viral-identify module is faster than alternative pipelines; There are currently, to our knowledge, no other peer reviewed softwares that assess metagenomic viral contig completeness and contamination (ViralQC [[25]](https://www.zotero.org/google-docs/?kz2AzS) has been recently released, but is not yet peer-reviewed and tested).

CheckV-PyHMMER-annotated vContigs then undergo a simple filtering procedure, removing any CheckV annotations with exclusion warnings (Supplementary Table 10). If CheckV annotations of genome quality is “Not-determined” (happens when CheckV doesn't detect any viral genes in the sequence or when the completeness prediction falls outside the defined ranges for other quality categories) or “Low-quality” (0-50% completeness), we allow the following contigs to pass through the filtering step:

I. If vContig has 0 geNomad viral hallmark genes***, it must have geNomad score >= 0.99
II. If vContig has 1-5 geNomad viral hallmark genes, it must have geNomad score >= 0.95
III. If vContig has > 5 geNomad viral hallmark genes, it must have geNomad score >= 0.95
IV. Regardless of the number of geNomad viral hallmark genes, if the geNomad Marker Enrichment Score is in the top 10th percentile of all sequences, it will also pass the filtering.

***geNomad viral hallmark genes are manually curated genes known to be associated with viral function and their presence strongly indicates a viral sequence.

We have two main reasons for performing a filtering step; Firstly that the geNomad viral hallmark database is more comprehensive than that of CheckV’s hallmark gene database and the existence of a geNomad hallmark highly likely denotes a viral contig [[7]](https://www.zotero.org/google-docs/?ZY7wss). ii) Secondly that, unlike CheckV, geNomad adopts a hybrid framework for identifying and annotating viruses, and is able to take into account both intrinsic sequence characteristics (Sequence Branch and the IGLOO encoder) and gene-based information (Marker Branch), allowing geNomad to identify shorter viral sequences with no hallmark genes accurately [[7]](https://www.zotero.org/google-docs/?ELHesp). The final output of this process is a viral Operation Taxonomic Units (vOTU) database in the form of a fasta file used for downstream analysis.

##### 1.1.5. vomix viral-taxonomy -h

After creating a vOTU database, vOMIX-MEGA determines the taxonomy of each sequence using geNomad (which is highly accurate and can determine up to Family level [[7]](https://www.zotero.org/google-docs/?s3zlTq)) and PhaGCN [[26]](https://www.zotero.org/google-docs/?USFLBW) (which can annotate up to species level, albeit the accuracy is yet to be tested by high quality benchmark data). We have also tested two other methods which we have decided to discard for the following reasons

1. NCBI-database approach implemented by Nayfach et. al pipeline [[17]](https://www.zotero.org/google-docs/?0vR4Fs): This method produced highly similar results to PhaGCN’s graph convolutional network model (Supplementary Table 11), likely since PhaGCN was trained on the same data. The PhaGCN model is pre-trained and hence much smaller in size and computationally less intensive than the NCBI alignment method, which is why we have chosen it over the NCBI method.
2. VIRify ViPhOG HMM-Approach [[27]](https://www.zotero.org/google-docs/?ndfSH5): This approach generated likewise similar results to geNomad’s classifications (Supplementary Table 11) but classified only 10% of what geNomad could classify. It is also a memory and computationally expensive procedure, which is why it was discarded in our pipeline.

The results of both geNomad and PhaGCN are compiled into one table for each vOTU and are available in a .csv file for further analysis.

##### 1.1.6. vomix viral-annotate -h

The viral annotate module takes input either the vOTUs generated from the `viral-identify` module or the user’s custom contig file to perform protein-level gene annotation. It generates gene annotation files using pyrodigal-gv [[28]](https://www.zotero.org/google-docs/?0NA2JV), and further annotates them using the eggNOG-mapper v2 [[29]](https://www.zotero.org/google-docs/?Xd8wAb), PhaVIP annotation [[9]](https://www.zotero.org/google-docs/?0tOIrE), MetaCerberus [[30]](https://www.zotero.org/google-docs/?qekBh2), and Pharokka [[31]](https://www.zotero.org/google-docs/?6Ofkms). It then merges these results into a concatenated .csv file called `viral_annotate_summary.csv`

##### 1.1.7. vomix viral-host -h

The host identification module takes vOTU database or any fasta file by the user and analyses the host by either CHERRY[[32]](https://www.zotero.org/google-docs/?mHpZV0) or iPHoP [[33]](https://www.zotero.org/google-docs/?3MOQFH). CHERRY is the default software, since it uses significantly less memory than iPHoP, as well as gives functional insight between shared genes of viruses and their host [[32]](https://www.zotero.org/google-docs/?B1H6EM). CHERRY likewise is able to show multi-host dynamicity using a network graph, which is a shortcoming of most viral host identification softwares that are only able to predict one host per virus [[34]](https://www.zotero.org/google-docs/?yD3gTB). iPHoP is memory intensive, time-consuming, hosts a very large database, and can identify one host per virus, but we have included it as an alternative as it uses interpretable alignment methods of CRISPR and non-CRISPR sequences to give a consensus identification of viruses and their host, allowing for possibly better biological interpretability [[33]](https://www.zotero.org/google-docs/?Pkq6HO).

##### 1.1.8. vomix viral-community -h

The viral-community module quantifies the abundance of vOTUs in the cleaned post-processed samples, and therefore needs a *sample_list.csv* file to point at the paired-end files. The module uses CoverM [[35]](https://www.zotero.org/google-docs/?DJ3m0L) contig mode to quantify the sequences and creates a Reads Per Kilobase per Million mapped reads (RPKM) and Transcripts Per Million mapped reads (TPM) vOTU abundance file.

##### 1.1.9. vomix viral-benchmark -h

The `viral-benchmark` module is built-in to vOMIX-MEGA to allow quick comparison of the outputs generated by 6 different viral contig annotation tools routinely used in viral genomics. It runs analysis on either a single fasta file or a group of fasta files in a directory treated as separate samples, and it analyses them with geNomad [[7]](https://www.zotero.org/google-docs/?rbBJDR), DeepVirFinder [[8]](https://www.zotero.org/google-docs/?hBBMKW), PhaMer [[9]](https://www.zotero.org/google-docs/?xAstu2), VirSorter2 [[10]](https://www.zotero.org/google-docs/?7blvEj), VirFinder [[11]](https://www.zotero.org/google-docs/?waKMue), and VIBRANT [[12]](https://www.zotero.org/google-docs/?sCV3cr). All outputs are summarised within one file called the `viral_benchmarking_summary.csv`.

##### 1.1.10. vomix prok-binning -h

All prokaryotic modules in vOMIX-MEGA are supplementary to the viral analysis and hence have standard and well-practiced protocols. The prokaryotic binning module takes contig from the assembly module and first maps the contigs to paired-end cleaned fastq files using strobealign [[36]](https://www.zotero.org/google-docs/?AfQkX4). It then either takes a multi-tool binning consensus approach using MetaBat2 [[37]](https://www.zotero.org/google-docs/?A1kA33), MaxBin [[38]](https://www.zotero.org/google-docs/?Gfcrh7), and CONCOT [[39]](https://www.zotero.org/google-docs/?zYL8QR), and refined using DasTool [[40]](https://www.zotero.org/google-docs/?1KPOgi). Alternatively, a modern GPU-based approach for binning can be used with VAMB if GPU processing is available to the user [[41]](https://www.zotero.org/google-docs/?GHLj4K). The final bins are then assessed for completion and contamination using CheckM2 [[42]](https://www.zotero.org/google-docs/?pNUYHM) and then dereplicated using galah [[43]](https://www.zotero.org/google-docs/?fFtKzJ). We then use GTDB-Tk v2 to quantify the taxonomy of each prokaryotic bin [[44]](https://www.zotero.org/google-docs/?wzc3X9).

##### 1.1.11. vomix prok-community -h

The *vomix prok-community module* is a Snakemake wrapper around the MetaPhlAn v4 [[45, p. 4]](https://www.zotero.org/google-docs/?24EWnr), and facilitates databases and environment installation and performs community quantification according to the MetaPhlAn protocol. The MetaPhlAn output is then stratified on taxonomy level and produces different abundance tables.

##### 1.1.12. vomix prok-annotate -h

The vomix prok-annotation is a simplified wrapper around HUMAnN3 analysis [[46, p. 3]](https://www.zotero.org/google-docs/?DScQlg). It performs HUMAnN3 analysis, then subsequently normalizes the results and regroups tables based on the Level-4 Enzyme Commission, EggNOG including COGs, Gene Ontology, KEGG Orthogroups, Pfam domains, and MetaCyc reactions outputs. It automatically creates both taxonomy-stratified and aggregated results.

##### 1.2. Iterative clustering algorithm

##### 1.2.1. Motivation and Conceptual Framework

In high-throughput viral metagenomics, clustering fragmented viral genomes to generate viral Operational Taxonomic Units (vOTUs) represents a severe computational bottleneck. Standard sequence-clustering algorithms—such as CD-HIT or CheckV’s native MEGABLAST-based pairwise alignment—typically require constructing exhaustive sequence databases or performing all-versus-all alignments. The memory footprint (Resident Set Size; RSS) of these indexing and comparison steps scales quadratically (
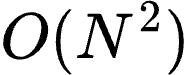
<https://saxarona.github.io/mathjax-viewer/?input=O(N%5E2)#0>) with the number of input sequences
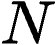
<https://saxarona.github.io/mathjax-viewer/?input=N#0>. When scaling to terabyte-scale metagenomic datasets, this mathematical scaling leads to massive memory inflation and execution hangs, frequently requiring computing architectures that are inaccessible to most researchers.

To bypass these hardware limitations, vOMIX-MEGA introduces an algorithmically re-engineered, loss-free data-splitting and iterative clustering framework (--splits). This module processes large nucleotide FASTA inputs in parallel, isolated fractions before performing hierarchical pooling and clustering. By integrating this partitioning logic with CheckV's MEGABLAST-based clustering mechanism, vOMIX-MEGA circumvents CD-HIT execution hangs on divergent datasets and maintains a stable memory footprint.

##### 1.2.2. Mathematical Formulation of Iterative Clustering

The iterative clustering algorithm is structured as a binary tree reduction (or tournament-style merger). The depth and branching factor of the tree are dynamically controlled by a user-defined parameter
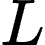
<https://saxarona.github.io/mathjax-viewer/?input=L#0>, representing the total clustering iterations (clustering_iter).

The system operates according to the following mathematical rules:

- Initial Partitioning: The raw input FASTA is divided into
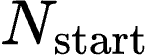
<https://saxarona.github.io/mathjax-viewer/?input=N_%7B%5Ctext%7Bstart%7D%7D#0> starting chunks:

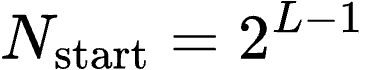
<https://saxarona.github.io/mathjax-viewer/?input=N_%7B%5Ctext%7Bstart%7D%7D%20%3D%202%5E%7BL-1%7D#0>
- Layer and Chunk Indexes: Let
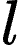
<https://saxarona.github.io/mathjax-viewer/?input=l#0> denote the current layer of the reduction tree, where
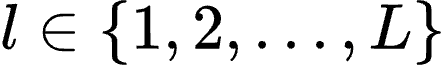
<https://saxarona.github.io/mathjax-viewer/?input=l%20%5Cin%20%5C%7B1%2C%202%2C%20%5Cdots%2C%20L%5C%7D#0>. Let
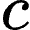
<https://saxarona.github.io/mathjax-viewer/?input=c#0> denote the chunk index within layer
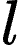
<https://saxarona.github.io/mathjax-viewer/?input=l#0>, where
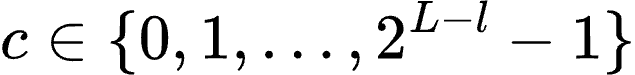
<https://saxarona.github.io/mathjax-viewer/?input=c%20%5Cin%20%5C%7B0%2C%201%2C%20%5Cdots%2C%202%5E%7BL-l%7D-1%5C%7D#0>.
- Base Split Inputs: Let
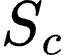
<https://saxarona.github.io/mathjax-viewer/?input=S_c#0> represent the
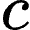
<https://saxarona.github.io/mathjax-viewer/?input=c#0>-th partitioned raw FASTA file.
- Clustering Operator: Let
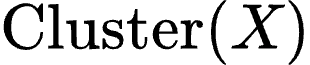
<https://saxarona.github.io/mathjax-viewer/?input=%5Ctext%7BCluster%7D(X)#0> denote the sequence clustering and dereplication function, executed using either CD-HIT or CheckV's MEGABLAST-based pairwise alignment.

The recursive generation of clustered outputs
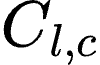
<https://saxarona.github.io/mathjax-viewer/?input=C_%7Bl%2C%20c%7D#0> at any given node of the tree is defined by the recurrence relation:

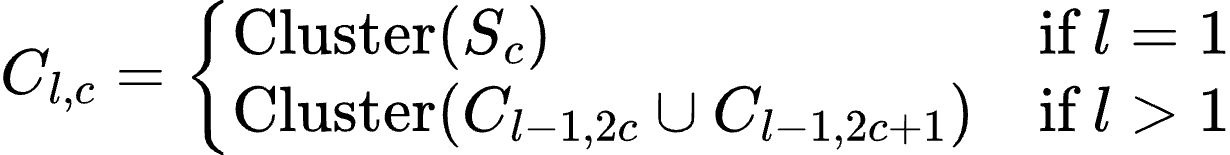
<https://saxarona.github.io/mathjax-viewer/?input=C_%7Bl%2C%20c%7D%20%3D%20%5Cbegin%7Bcases%7D%20%5Ctext%7BCluster%7D(S_c)%20%26%20%5Ctext%7Bif%20%7D%20l%20%3D%201%20%5C%5C%20%5Ctext%7BCluster%7D(C_%7Bl-1%2C%202c%7D%20%5Ccup%20C_%7Bl-1%2C%202c%2B1%7D)%20%26%20%5Ctext%7Bif%20%7D%20l%20%3E%201%20%5Cend%7Bcases%7D#0>

Using this binary tree architecture, the total number of individual clustering executions (
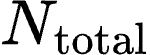
<https://saxarona.github.io/mathjax-viewer/?input=N_%7B%5Ctext%7Btotal%7D%7D#0>) required to resolve the complete dataset down to a single, fully non-redundant output chunk (
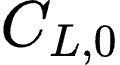
<https://saxarona.github.io/mathjax-viewer/?input=C_%7BL%2C%200%7D#0>) is given by:

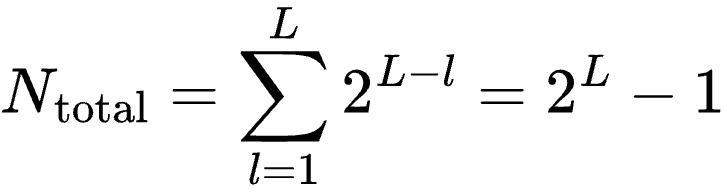
<https://saxarona.github.io/mathjax-viewer/?input=N_%7B%5Ctext%7Btotal%7D%7D%20%3D%20%5Csum_%7Bl%3D1%7D%5E%7BL%7D%202%5E%7BL-l%7D%20%3D%202%5EL%20-%201#0>

##### 1.2.3. Comparative Memory Complexity: Brute Force vs. Tournament Reductions

To formally demonstrate why the recursive approach utilizes less memory, we contrast its mathematical peak memory bounds against the standard brute-force method.

##### 1.2.3.1. The Brute Force Approach (No Partitioning)

If all
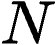
<https://saxarona.github.io/mathjax-viewer/?input=N#0> sequences are clustered or indexed in a single pass (as in conventional, non-optimized workflows), peak memory consumption is an unconstrained function of the total raw input size:

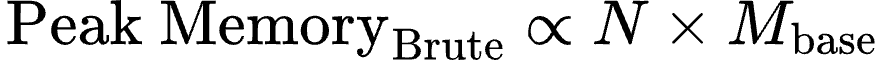
<https://saxarona.github.io/mathjax-viewer/?input=%5Ctext%7BPeak%5C%20Memory%7D_%7B%5Ctext%7BBrute%7D%7D%20%5Cpropto%20N%20%5Ctimes%20M_%7B%5Ctext%7Bbase%7D%7D#0>

where
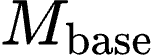
<https://saxarona.github.io/mathjax-viewer/?input=M_%7B%5Ctext%7Bbase%7D%7D#0> represents the baseline memory footprint (indexing and overhead metadata) required per sequence.

##### 1.2.3.2. The Tournament Protocol (Partitioning + Dereplication)

Under vOMIX-MEGA’s --splits protocol, the sequence dataset is partitioned into
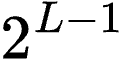
<https://saxarona.github.io/mathjax-viewer/?input=2%5E%7BL-1%7D#0> initial chunks and progressively collapsed. Let
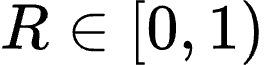
<https://saxarona.github.io/mathjax-viewer/?input=R%20%5Cin%20%5B0%2C%201)#0> represent the dataset's redundancy factor, which denotes the fraction of redundant sequences collapsed during clustering. The peak memory footprint of this protocol is strictly bounded at two distinct phases:

- **Phase 1** (Base Layer Memory): At the initial split layer (
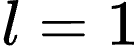
<https://saxarona.github.io/mathjax-viewer/?input=l%20%3D%201#0>), the data is divided into
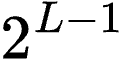
<https://saxarona.github.io/mathjax-viewer/?input=2%5E%7BL-1%7D#0> chunks. The peak memory required to cluster any individual chunk is scaled down to:

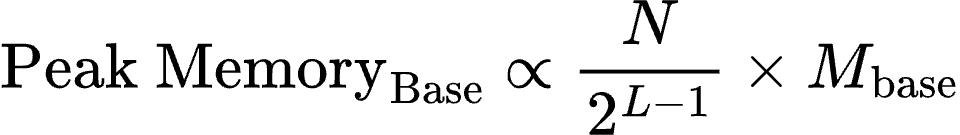
<https://saxarona.github.io/mathjax-viewer/?input=%5Ctext%7BPeak%5C%20Memory%7D_%7B%5Ctext%7BBase%7D%7D%20%5Cpropto%20%5Cfrac%7BN%7D%7B2%5E%7BL-1%7D%7D%20%5Ctimes%20M_%7B%5Ctext%7Bbase%7D%7D#0>

- **Phase 2** (Upper Layer Memory): As merged files propagate up the binary tree, redundancy is filtered out. By the final iterations, the unique sequence pool has shrunk, bounding the memory footprint of the final merge step to:

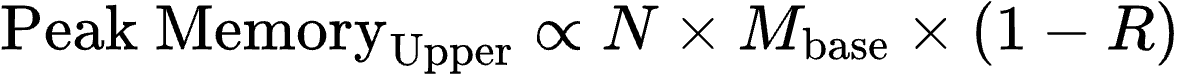
<https://saxarona.github.io/mathjax-viewer/?input=%5Ctext%7BPeak%5C%20Memory%7D_%7B%5Ctext%7BUpper%7D%7D%20%5Cpropto%20N%20%5Ctimes%20M_%7B%5Ctext%7Bbase%7D%7D%20%5Ctimes%20(1%20-%20R)#0>

##### 1.2.3.3. Synthesis and mathematical guarantee of memory reduction

The global peak memory footprint of the tournament-style iterative clustering pipeline is determined by the maximum resource bottleneck across these two phases:

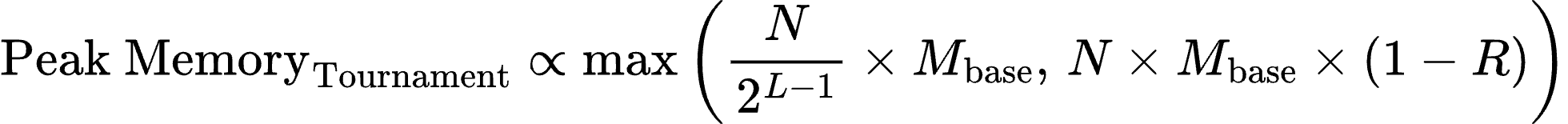
<https://saxarona.github.io/mathjax-viewer/?input=%5Ctext%7BPeak%5C%20Memory%7D_%7B%5Ctext%7BTournament%7D%7D%20%5Cpropto%20%5Cmax%5Cleft(%20%5Cfrac%7BN%7D%7B2%5E%7BL-1%7D%7D%20%5Ctimes%20M_%7B%5Ctext%7Bbase%7D%7D%2C%5C%2C%20N%20%5Ctimes%20M_%7B%5Ctext%7Bbase%7D%7D%20%5Ctimes%20(1%20-%20R)%20%5Cright)#0>

Because
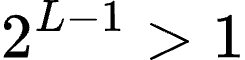
<https://saxarona.github.io/mathjax-viewer/?input=2%5E%7BL-1%7D%20%3E%201#0> (for any iteration depth
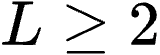
<https://saxarona.github.io/mathjax-viewer/?input=L%20%5Cge%202#0>) and metagenomic sequence cohorts typically possess a high sequence redundancy factor (
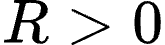
<https://saxarona.github.io/mathjax-viewer/?input=R%20%3E%200#0>), the following inequality holds:

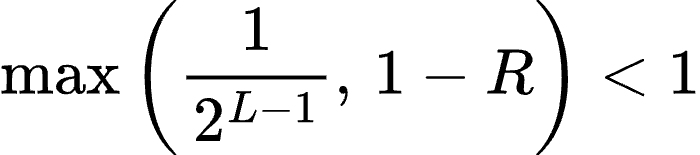
<https://saxarona.github.io/mathjax-viewer/?input=%5Cmax%5Cleft(%5Cfrac%7B1%7D%7B2%5E%7BL-1%7D%7D%2C%5C%2C%201-R%5Cright)%20%3C%201#0>

Consequently:

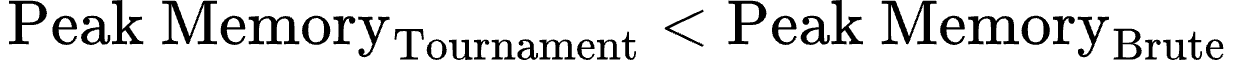
<https://saxarona.github.io/mathjax-viewer/?input=%5Ctext%7BPeak%5C%20Memory%7D_%7B%5Ctext%7BTournament%7D%7D%20%3C%20%5Ctext%7BPeak%5C%20Memory%7D_%7B%5Ctext%7BBrute%7D%7D#0>

This inequality formally proves why vOMIX-MEGA maintains a highly constrained, stable memory footprint even when scaling to terabyte-scale inputs.

##### 1.3. vOMIX-MEGA Input Formats

vOMIX-MEGA takes multiple flexible input formats depending on the module being used. This allows full customization for users to use each step of the pipeline they find suitable. The main inputs are:

1. **sample_list.csv**: A comma-separated table with either SRA accessions, sample IDs and optionally R1 and R2 paired read files locally stored on the user’s computer. Used when needing to read paired-end files as input to a module. The *sample_list.csv* file allows both local and remote files to be used as input into the user’s analysis. SRA samples are automatically downloaded and local files are validated.
2. **fasta file**: A single fasta nucleotide file ending with .fa or .fasta extensions. Used for modules that only require a list of contigs or fasta sequences as inputs. For example, *vomix clustering-fast* module can take one concatenated list of contigs that it will cluster according to specified thresholds.
3. **fasta directory:** A directory with fasta files with the .fa or .fasta files. Used for modules that can take multiple separated fasta files as inputs and treated as separate samples. For example, *vomix viral-identify* module where a directory of different contig files from different samples can be used as input and each individual fasta file will be treated as a separate sample.

You can read more about the input types in our Wiki page <https://vomix-mega.readthedocs.io/en/latest/>.

[[46]](https://www.zotero.org/google-docs/?ZucSTl)	F. Beghini *et al.*, “Integrating taxonomic, functional, and strain-level profiling of diverse microbial communities with bioBakery 3,” *eLife*, vol. 10, p. e65088, doi: 10.7554/eLife.65088.
